# Estimation of the lag time in a subsequent monomer addition model for fibril elongation

**DOI:** 10.1101/034900

**Authors:** Suzanne K. Shoffner, Santiago Schnell

## Abstract

Fibrillogenesis, the production or development of protein fibers, has been linked to protein folding diseases. The progress curve of fibrils or aggregates typically takes on a sigmoidal shape witha lag phase, a rapid growth phase, and a final plateau regime. The study of the lag phase and the estimation of its critical timescale provide insight into the factors regulating the fibrillation process. However, methods to estimate a quantitative expression for the lag time rely on empirical expressions, which cannot connect the lag time to kinetic parameters associated with the reaction mechanisms of protein fibrillation. Here we introduce an approach for the estimation of the lag time using the governing rate equations of the elementary reactions of a subsequent monomer addition model for protein fibrillation as a case study. We show that the lag time is given by the sum of the critical timescales for each fibril intermediate in the subsequent monomer addition mechanism and therefore reveals causal connectivity between intermediate species. Furthermore, we find that single-molecule assays of protein fibrillation can exhibit a lag phase without a nucleation process, while dyes and extrinsic fluorescent probe bulk assays of protein fibrillation do not exhibit an observable lag time phase during template-dependent elongation. Our approach could be valuable for investigating the effects of intrinsic and extrinsic factors to the protein fibrillation reaction mechanism and provides physicochemical insights into parameters regulating the lag phase.

## 1 Introduction

Protein folding is vital to normal functioning of the cell. While most proteins have one or more native conformations, some proteins misfold into a non-native conformation, causing accumulation and ultimately the formation of amorphous aggregates, or in the case of amyloidogenic proteins, mature amyloid fibrils.^1,2^ Fibrils generally provide a more stable conformation than the anomalous state due to the stabilization of cross-*β*-sheets by the peptide backbone.^3^ Though the formation of dimers, trimers, and other larger oligomeric complexes are part of normal, healthy cell functioning, aberrant protein aggregates can be toxic and have been shown to have pathological consequences.^4^ Protein aggregation has been linked with over 50 protein folding diseases and disorders, including type II diabetes and Alzheimer’s, Parkinson’s, and Huntington’s disease.^2^ Though significant progress has been made toward understanding the reaction mechanisms of protein aggregation for some diseases (i.e. Alzheimer’s disease via A*β* propagation^5^), the large majority of proteins aggregate with mechanisms that remain to be identified. Fibrillogenesis is a complex multistep process, generally beginning with monomers or other small molecules that collide and bond to form larger molecules including oligomers and protofibrils, until the fibril sizes formed have reached equilibrium.^6^ The equilibrium fibril size distribution can vary from strongly skewed distributions to broad distributions depending on a number of factors, including elapsed time, fragmentation effects, and aggregate merging.^7,8^ Estimation of the rates of aggregation reactions will be important not only for identification and overall understanding of aggregation mechanisms, but also for developing pharmacological treatment and strategies for disease prevention.

In a typical protein aggregation kinetic experiment, the time course of protein fibrillation is measured by the absorbance of light at one or more wavelengths using dyes and extrinsic fluorescent probes.^9^ Time course data often follow the characteristic shape of a sigmoidal curve and are separated into three regions: a lag phase, a fast growth phase, and a plateau phase. The lag phase is of particular interest because it provides critical information about the factors regulating the fibrillation process. A major challenge is determining which molecular events regulate the lag phase in fibril formation.^10^ There has been much debate on whether nucleation, growth, or both contribute to the lag phase of the sigmoidal-shaped curve.^11,12^ Another important point to take into consideration is that the interpretation of the lag time is dependent on the experimental technique used to make kinetic measurements. For example, the fluorescence dye Thioflavin T (ThT) is used to probe protein aggregation,^13^ because it binds to *β*-sheet-rich structures and does not affect the aggregation kinetics. ^14^ The formation of aggregates can be probed by monitoring the increase in the fluorescence as a function of time in a multiwell plate reader format. ThT fluorescence intensity is not an absolute measure of the amount of protein fibrils formed because multiple ThT molecules bind to multiple species, including prefibrillar oligomers and fibrils of differing sizes. Therefore, the lag time of a ThT assay does not correspond to a particular molecular event and cannot be attributed to primary nucleation events alone.^15^ In contrast, single-molecule studies are used to monitor the time course of individual fibril species, and thus, a lag time for each fibril of different size can be potentially estimated. New single-molecule techniques are rapidly emerging. For example, Horrocks et al.^16^ developed a technique called Single Aggregate Visualization by Enhancement which allows for ultra-sensitive fluorescent detection of individual amyloid fibrils. Tosatto et al.^17^ found that single-molecule Förster Resonance Energy Transfer helps to visualize the self-assembly process using one-channeled microfluidic devices.^18^ Pinotsi et. al^19,20^ used superresolution optical nanoscopy to examine the nucleation and elongation processes on a molecular level.

In both the dyes/extrinsic fluorescent probe bulk assays and sophisticated single-molecule assays, the lag phase is typically studied with an empirical logistic (sigmoidal) function, which is used to estimate phenomenological parameters from fibrillation time course data. The basic logistic function gives the characteristic sigmoidal shape, but is limited to describing symmetrical progress curves.^21^ The Gompertz function and the Richards function exhibit the asymmetrical sigmoidal shape commonly observed with fibrillogenesis. ^22,23^ The Richards function adds an additional parameter, *v*, to account for asymmetry:

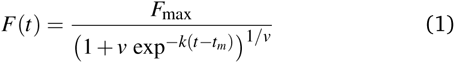

where *F*(*t*) is the fluorescence intensity at time *t*, *F*_max_ is the steady-state fluorescence at the plateau of the progress curve, *t*_*m*_ is the inflection time at which the growth rate reaches its maximum (*v*_max_), and *v* is a parameter that alters the asymmetry of the curve. A geometric representation of these parameters is shown in **Fig. 1**. Eqn (1) and variations thereof are often used to estimate the empirical parameters for progress curves of protein aggregation experiments.^24–26^

**Fig. 1.**
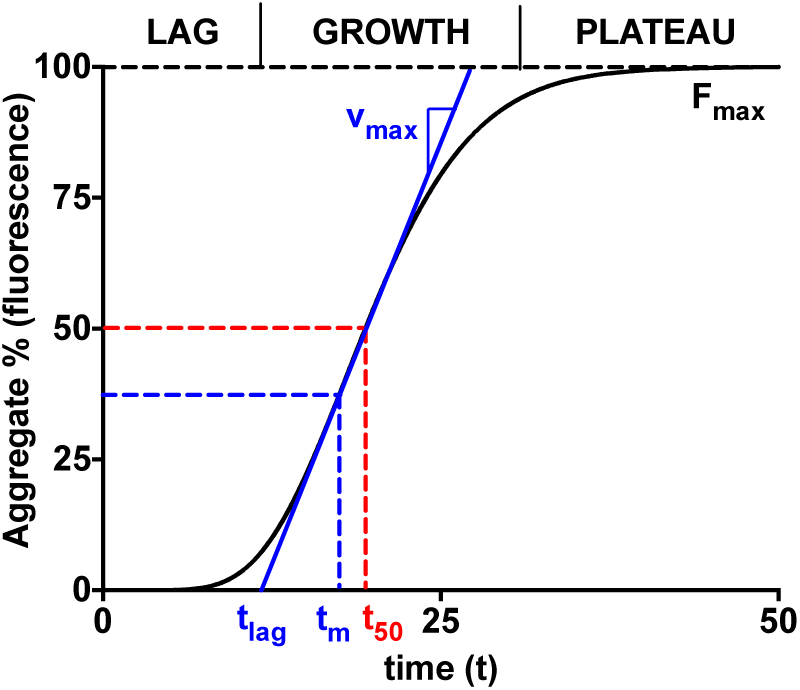
The increasing concentration of aggregates as a function of time displays the characteristic sigmoidal curve for amyloid fibril formation. Aggregate concentration is represented as fluorescent intensity percentage of the final aggregates (*F*_max_). The curve is typically divided into the lag phase, the growth phase, and the plateau phase. The half-time *t*_50_ is the time at which half of the plateau aggregates are formed. The inflection time *t*_*m*_ is when the growth rate reaches its maximum, *v*_*max*_. The lag time is then typically estimated by extending the tangent at *t*_*m*_ down to the time axis.

The lag time (*t*_lag_) is typically estimated by extending the tangent at *t*_*m*_ to the initial baseline. The lag time is then given in terms of the empirical parameters:

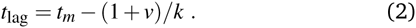

Another type of critical timescale is the amount of time needed for the percentage of aggregate formed to reach a certain threshold. For example, *t*_50_ (*t*_1/2_), often called the halftime, refers to the amount of time required to reach half of the maximum fluorescence intensity or percentage of the final aggregate. Occasionally, researchers will choose an arbitrary value, such as 10%, for the threshold. However, having multiple definitions for critical lag times introduces variability and uncertainty in estimates reported in the literature. At the same time, phenomenological time lag estimates are unable to provide a relationship between the model parameters and the elementary reaction steps governing the reaction mechanisms.

In amyloid studies, samples can involve billions of monomers and the reaction schemes may be very complex.^10^ There are many possibilities for fragmentation and association when considering the reversible association of polymers.^27^ Polymer fragmentation may also lead to secondary nucleation, since fragmented polymers may serve as nuclei for elongation.^15^ Fragmentation and other molecular events, such as inhibition, and off-pathway aggregation, have been predicted to contribute significantly to the lag phase of fibril formation. For example, Pagano et al.^25^ show that targeting the initial steps of fibrillation via kinetic inhibition reduces the concentration of early intermediate size oligomers. The onset of fibrillation relates to the concentration of unbound protein species in the presence of an inhibitor. In another study by Liu et al.^26^ the lag time for A*β*40 fibrillogenesis is slightly elongated in the presence of a protein aggregation inhibitor, chitosan. Inhibition kinetics may therefore be particularly relevant for research on aggregation prevention strategies. The lag time may also be affected by off-pathway aggregation that leads to the formation of toxic deposits. For example, a study by Crespo et al.^8^ suggests that competitive off-pathway steps may be favored for monomer addition and therefore increase the lag time for fibrillation. Additionally, environmental factors such as pH and temperature can have an effect on the shape of the curve and the resulting estimated lag time.^28^

Reaction kinetic and thermodynamic models have sought to describe the process mechanistically. They range in complexity, from simpler subsequent monomer addition models to complex nucleation and elongation models.^29,30^ In this paper, we propose a protocol for deriving an expression for the lag time from the governing rates of a nucleation-independent, subsequent monomer addition model. The goal is to find a mathematical expression for the lag time in terms of the reaction parameters that is based on reaction kinetics rather than on empirical sigmoidal equations. We introduce an expression for the observable formation of fibrils to experimentally validate our model and explore the differences between the lag time in single-molecule and bulk analytical assays. By focusing on the elongation and growth stage of the process, we are able to determine whether growth alone is sufficient to produce the characteristic sigmoidal shape for the time course of a single molecule fibril species. This work introduces a novel approach to derive an expression for the lag time that may be more meaningful molecularly to the protein aggregation community, while allowing us to introduce the underlying molecular details of a subsequent monomer addition mechanism.

## 2 A subsequent monomer addition dock-lock reaction mechanism for fibril elongation

Since the initiation process by which native monomers misfold and nucleate is complex and largely based on random events, in this paper, we focus on the highly organized elongation of the fibrils. Our model is based on the Esler et al.^5^ dock-lock growth mechanism, where the monomer *M* first reversibly “docks" to an initial fibril template *F*_0_, creating a monomer-template complex which we call *C*_1_. After the rapid “docking" stage, the “docked" monomer undergoes a slow conformational change and becomes irreversibly associated with the template, forming the “locked" fibril *F*_1_. The “locked" fibril goes on to serve as a template for the deposition of another monomer, such that the most recently deposited monomer dissociates first, while previously deposited monomers become “locked" onto the template. The “locked" fibrils are denoted by *F*_*i*_ (*i* = 1…*n*, where *i* indicates the number of “locked" monomers added to the template) undergo elongation via subsequent monomer addition until the final fibril (*F*_*n*_) is synthesized. The reaction scheme of this fibril elongation reaction mechanism is given by

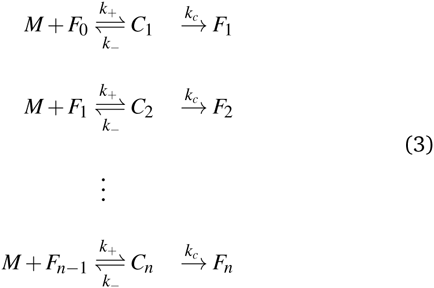

where *k*_+_ and *k*_–_ are the on‐ and off-rate constants, respectively. The synthesis rate for the “locked" fibrils is given by *k*_*c*_. Applying the law of mass action to reaction scheme (3) yields a nonlinear system of 2*n* + 2 ordinary differential equations of the form:

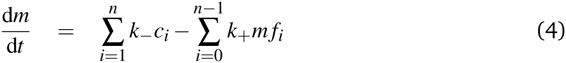

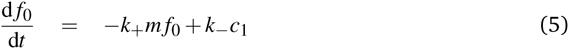

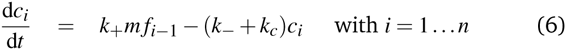

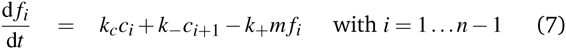

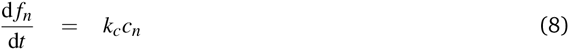

where the lowercase indicates the concentration of that species. The initial conditions for the reaction scheme are given by 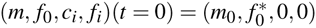 with *i*= 1… *n*.

The above system has two conservation laws: the total fibril and total monomer are conserved in the free (in solution) and bound (“docked” or “locked”) state. Assuming that the reaction is a closed and isolated thermodynamic system, we obtain a mass conservation law for the total fibril, given by a sum of the fibril template *f*_0_ and the “docked" *c*_*i*_ and “locked" fibril *f*_*i*_:

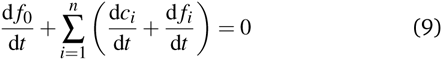

which implies that

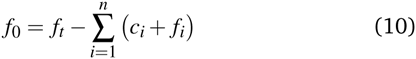

where *f*_*t*_ is the total concentration of fibril. Given that the monomer is also carried through the process in the free (in solution) or bound (“docked” or “locked”) state, there is a second conservation law for the monomer in solution *m* and the “docked” *c*_*i*_ and “locked” monomer *f*_*i*_:

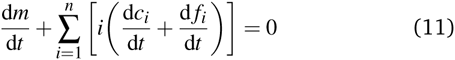

which implies that

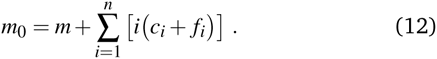

We can use these conserved quantities to simplify the ordinary differential equation system, thereby reducing its dimension. We combine the rate constants into physically meaningful parameters, where 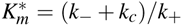 is the apparent dissociation constant of the monomer from the “docked” monomer-template complex, *K*_*m*_ = *k*_–_/*k*_+_ is the dissociation constant of the monomer from the “docked” monomer-template complex, and *K* = *k*_*c*_/*k*_+_ is the fibrillation constant. ^31^

## 3 Estimation of the fibrillation time lag

From the physicochemical point of view, the intermediate “docked" monomer-template complex species are short-lived and react quickly during an initial transient, *t*_*c*_, of reaction mechanism (3). During this period, we assume (**Assumption I**) that the concentration of the monomer (*m*) and the concentration of the fibril template (*f*_0_) remain approximately constant. For *t* < *t*_*c*_,

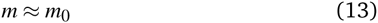

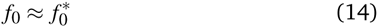

This assumption is known as the reactant stationary approximation,^32^ and the conditions for its validity are presented in Section 4.1. Using the reactant stationary approximation, we can estimate the timescale for a significant increase in the concentration of the “locked" fibrils (*t*_*f*_*i*__) using a mathematical scaling and simplification technique similar to Segel.^33^ However, we first need to derive a solution for the time course of the “docked" monomer-template complex during the initial transient (*t* < *t*_*c*_). We substitute eqn (13) into eqn (6) and obtain:

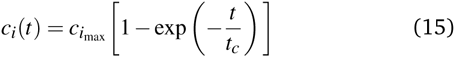

where *c*_*i*_*max*__ is the maximum concentration of the intermediate “docked" monomer-template complex *i* during the reaction

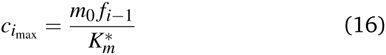

and *t_*c*_* is the critical timescale of the intermediate “docked" monomer-template complexes

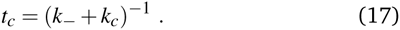

After the initial transient (*t* > *t*_*c*_), the “docked" monomer-template complexes and the monomer start to be depleted. As the reaction progresses, the concentrations of each intermediate “docked" monomer-template complex become significantly smaller than the initial monomer concentration (*m*_0_), such that

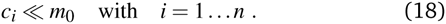

By applying eqn (18) to the conservation law for the monomer eqn (12), we find an expression for the concentration of the monomer after the initial transient

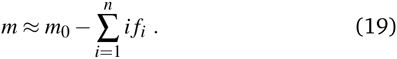

We can then approximate the depletion of the monomer at the timescale for each “locked" fibril (*t*_*f*_*i*__) using the reaction stoichiometry. We assume (**Assumption II**) that the fibril template *F*_0_ is the limiting reactant and that the monomer is in excess. Since there is a stoichiometric ratio of 1:1 between the reactant *F*_0_ and the first “locked" fibril product *F*_1_, we know that the maximum number of molecules of *F*_1_ produced will be equal to the the initial number of fibril template molecules if the fibril template is the limiting reactant. In the next step of the reaction, the monomer is again assumed to be in excess, and the maximum number of *F*_2_ molecules produced from the limiting reactant *F*_1_ will again be equal to the initial number of fibril template molecules. This will continue for each step of the reaction, such that *m* is depleted by approximately the concentration of 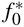 for each step. Therefore, the approximate final steady-state concentration for the monomer will be equal to

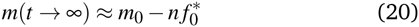

and *m* at the timescale for each “locked" fibril *t*_*f*_*i*__ can be approximated by the following expression:

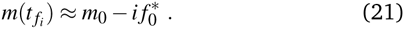

We then substituted eqn (15) into eqn (7), using eqn (21) for the depletion of *m*. The time course for the fibril after the intermediate “docked" monomer-template complex reaches its maximum is then given by

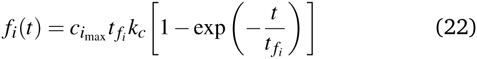

where *t*_*f*_*i*__ is the timescale of the “locked" fibrils

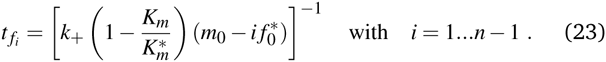

The timescales for each species (*t*_*c*_ and *t*_*f*_) are defined as the absolute value of the reciprocal of the eigenvalue for that species.^32^ They are only dependent on kinetic parameters of the reaction mechanism and the initial concentration of monomer or initial fibril template. The underlying physicochemical principle behind **Assumption II** is that each preceding intermediate “locked” fibril must form before the subsequent “locked" fibril size is formed. Given that the intermediate “locked” fibrils are ultimately depleted and the monomer is in excess, we can then estimate the critical lag time for significant production of the final fibril *f*_*n*_ by summing the timescales for all preceding “locked” fibril intermediates:

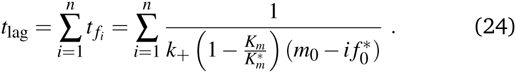

The lag time describes causal connectivity of the “locked” intermediate fibrils in the reaction mechanism, showing that as the number of reaction steps between the final product and the initially perturbed species (the monomer and initial fibril template species) increase, the separation time between the maxima of the intermediate “locked” fibrils increase. The subsequent monomer addition can be compared to a set of falling dominos; after an initial perturbation of the reactant, each subsequent species must wait for the previous species to form (or fall). The waiting time for each domino, given that the previous domino has just begun to fall, is analogous to the timescale of each intermediate “locked” fibril *t*_*f*_*i*__. The lag time, equal to the total time before the last domino falls, is the sum of the waiting times for every domino, analogous to a significant formation of fibril *f*_*n*_ at *t*_lag_. Upon plotting the intermediate “locked’ fibrils, the ’domino effect’ is observed, showing that the lag time is equal to the sum of the timescales required for each previous “locked” fibril to form. The estimated lag time appears similar to a halftime *t*_50_ definition, though it has a direct relationship with the physicochemistry and kinetics parameters of the reaction mechanism (3). The timescales for each “locked” fibril, calculated using eqn (23), and the lag time for the final fibril, calculated using eqn (24), are shown in **Fig. 2**. The system of ordinary differential equations was solved using a variable order stiff differential equation solver in MATLAB^®^.

**Fig. 2.**
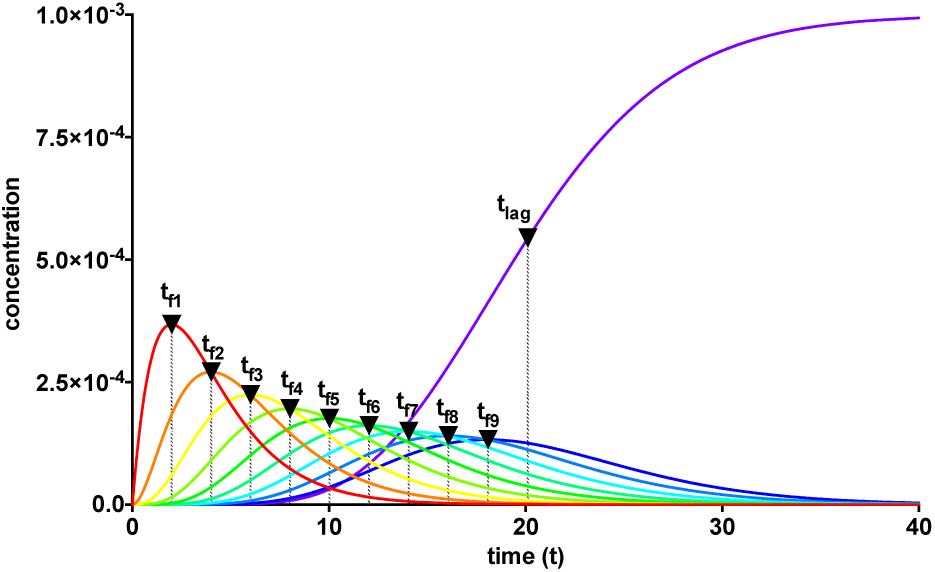
The lag phase for fibril elongation is characterized by the cumulative sum of the timescales for each of the intermediate “locked” fibrils, analogous to causal connectivity of falling dominos. Parameter values are: 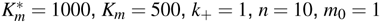 and 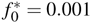

## 4 Conditions for validity of the lag time expression

In deriving the lag time, eqn (24), we made two assumptions: **Assumption I.** During the initial transient, *t* < *t*_*c*_, the reactants (monomer and fibril template) follow the reactant stationary approximation.^32^ **Assumption II.** After the initial transient, *t* > *t*_*c*_, the initial fibril template is the limiting reactant and the monomer is in excess. Under what experimental conditions are these assumptions valid?

### 4.1 Reactant stationary approximation

The reactant stationary approximation states that the depletion of the reactant is negligible during the initial transient, and therefore, the reactant concentrations will remain approximately constant during this time. ^32^ We can mathematically derive expressions for *m* ≈ *m*_0_ and 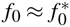 during the initial transient by writing that the decrease in reactant concentration (Δ*m* and Δ*f*_0_) is less than the product of the timescale of the initial transient (*t*_*c*_) and the initial maximal rate of reactant depletion.^33^ For the monomer, this expression is:

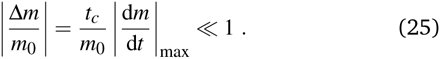

Substituting in eqn (17) and the maximal depletion rate from eqn (4), we find that the reactant stationary approximation for the monomer is given by the following inequality:

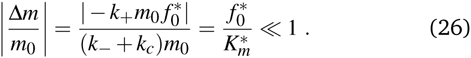

Similarly, we assume that the initial fibril template *f*_0_ remains approximately constant during the initial transient (*t* < *t*_*c*_):

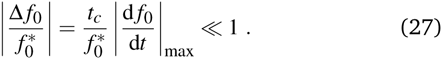

Substituting in eqn (17) and the maximal depletion rate from eqn (5), we obtain the reactant stationary approximation for the initial fibril template:

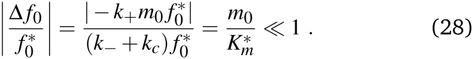

The reaction must satisfy the reactant stationary approximation conditions for the monomer and the initial fibril template, eqns (26) and (28), in order for the estimation of the lag time to be valid. Numerical confirmation for the validity of this approximation is shown in **Fig. 3**. We guarantee that eqn (26) and eqn (28) were met for all simulations, such that 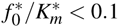 and 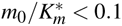

**Fig. 3.**
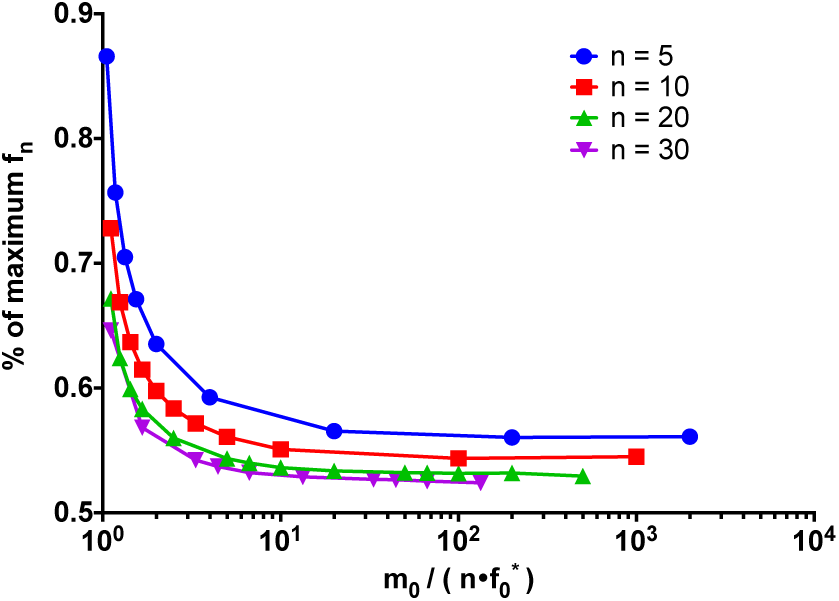
Numerical confirmation of the conditions for validity of the time lag expression. The reactant stationary approximation is always met, such that 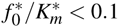 and 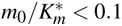. For the excess monomer condition 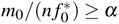 the variation in the percent of the maximum reached at *t*_lag_ is small. For *α* > 10^1^, the percentage is approximately 55%. The percentage of the maximum reached decreases at a decreasing rate as the length of the fibril *n* increases.

### 4.2 Monomer in excess and fibril template as the limiting reactant

Under **Assumption II**, we consider that the initial template fibril is the limiting reactant and the monomer is in excess during the time course of the reaction. The excess monomer condition ensures that there is stoichiometrically a sufficient amount of monomer for the reaction to finish to completion. The maximum amount of “locked” fibril created from the first step is stoichiometrically equal to the amount of limiting reagent. If the monomer is the limiting reagent, the intermediate “locked” fibrils are formed but not depleted. In the particular case we are investigating, intermediate “locked” fibrils are short-lived and ultimately depleted. To guarantee that this happens, we assume that the monomer is always in excess, taken from eqn (20), such that 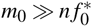 Then to test the strength of this condition, we add a factor *α* to describe the order of magnitude difference necessary to have stable conditions

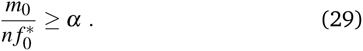

Eqn (29) allows us to test the validity of the lag time expression under different conditions of the ratio of reactant in excess to limiting reactant (*m*_0_ to 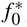), given that the conditions, eqns (26) and (28), for the reactant stationary approximation are met. With *m*_0_ fixed to one and the length *n* fixed, for large values of 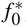, such that *α* < 10, the percentage of the maximum product reached at the lag time increases very rapidly and the excess monomer condition is unstable. When 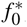 is small such that *α* > 10, the condition is very strong and there is very little variation in the lag time with changes to the ratio 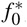/*m*_0_. Additionally, when the monomer is greatly in excess (more than one order of magnitude larger), the percentage of the maximum concentration of *f*_*n*_ approaches a constant value of approximately 55%. Therefore, when the excess monomer condition is met such that *m*_0_/(*nf*_0_) ≥ *α* for *α* > 10, the percentage of the maximum concentration of *f*_*n*_ reached approaches a constant for each size *n*. As *n* increases, the percentage of the maximum reached at *t*_*lag*_ approaches 50%, as shown in **Fig. 3**. This is further clarified upon further examination of eqn (24): when 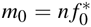 the denominator is zero and the lag time approaches infinity. If 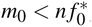 the term in the denominator becomes negative, and therefore, while the rate equations are still valid, our definition of the lag time is invalid in this region (see ESI **Fig. S1**).

## 5 Factors affecting the length of the lag time

Now we focus our attention to reaction kinetic factors influencing the length of the lag phase by systematically varying each of the parameters in eqn (24).

The elongation and size of the longest fibril is determined by *n*. As expected, the longer the fibril, the longer it takes to form. The lag time increases linearly with fibril length, as shown in **Fig. 4A**. For computational purposes, we selected small fibril sizes (*n* = 10,20,30,40,50), though fibrils can be composed of hundreds of thousands of monomers.

**Fig. 4.**
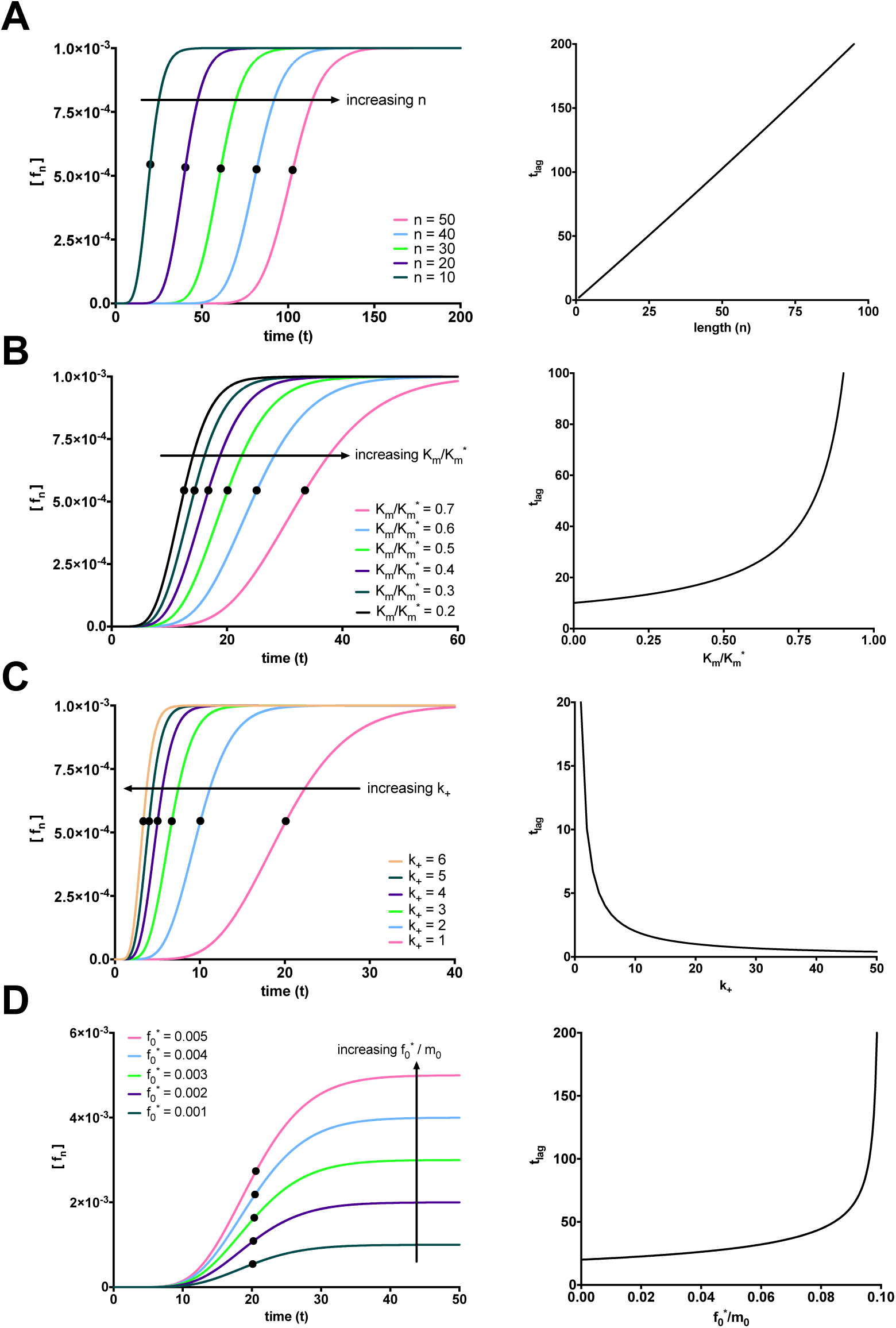
Factors affecting the length of the fibrillation lag time. (A) The time lag increases linearly with increasing fibril length. (Parameter values: 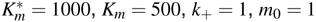, and 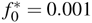). (B) The time lag increases exponentially with increasing ratio of *K*_*m*_ to 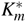 (Parameter values: 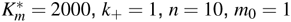 and *f*_0_ = 0.001). (C) The time lag decreases exponentially with increasing *k*_+_. (Parameter values: 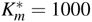, *K*_*m*_ = 500, *n* = 10, *m*_0_ = 1, and 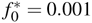). (D) Increasing the ratio of 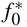 to *m*_0_ increases the plateau for the final fibril species (*f*_n_) and has little effect on the lag time in the range of 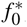 required for the condition of validity for eqn (29). (Parameter values: 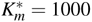, *K*_*m*_ = 500, *k*_+_ = 1, *n* = 10 and *m*_0_ = 1).

Changes to the kinetic constants also have an effect on the lag time. When examining how the ratio of the dissociation constant of the monomer from the intermediate “docked” monomer-template complex *K*_*m*_ to the apparent dissociation constant of the monomer from the intermediate “docked” monomer-template complex 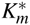 alters the lag time, we considered how much of an effect the monomer-fibril association-dissocation reaction has with respect to the entire reaction mechanism. Increases to 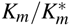 lead to an exponential increase in the lag time. When the overall reaction mechanism is shifted towards monomer-fibril “docked” complex association-dissociation, the reaction mechanism is shifted left and will take longer to form the longest fibril. The lag time exhibits an exponential increase with increasing 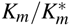 as shown in **Fig. 4B**.

The on-rate constant (*k*_+_) also affects the length of the lag time. An increase in the constant for association between a monomer and fibril (*k*_+_) shifts the overall reaction to the right and ultimately promotes fibril formation. The lag time exhibits an exponential decrease with increasing *k*_+_, as shown in **Fig. 4C**. We now turn our attention to the initial concentration of fibril template 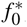 and the initial concentration of monomer mo. Note that increasing the ratio of 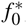 to *m*_0_ can violate the condition for monomer in excess and the fibril template as the limiting reactant eqn (29). Therefore, we can only increase 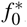 within a range where this condition is not violated. In this range, according to the conservation law for the total fibril, eqn (10), 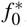 = *f*_*t*_ and *f*_*n*_(*t* →∞)= *f*_*t*_. The final fibril *f*_*n*_ approaches the initial concentration of the limiting reactant, the fibril template 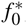. Increasing 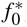 directly leads to a proportional increase in the plateau for the final fibril. The increase in *F*_*max*_ with increasing 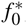/*m*_0_ is illustrated in **Fig. 4D**. When the excess monomer condition is met, there is a negligible, yet slight positive correlation between the lag time and the initial ratio of fibril template to monomer. If 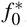 is fixed and mo (in excess) is increased, the lag time decreases exponentially (see ESI **Fig. S2**).

## 6 Experimental validation of fibril rate expressions as a function of time

The rate equations and expression for estimating the lag time described in this work can be applied directly to single-molecule studies which target the measurement of individual fibril species. As long as the reactant stationary approximation and monomer in excess conditions are met, the expression of the lag time will provide a relationship between 55% of the maximum concentration of the “locked” fibril and the kinetic parameters associated to the elementary steps governing for the formation of the fibrils. Currently, single-molecule studies that monitor changes in concentration or number of single fibril species as a function of time are limited in number. However, this situation is rapidly changing with the emergence of sensors and nanotechnology, and novel applications of single-molecule Förster Resonance Energy Transfer to directly visualize protein aggregation and fibrillation.

To experimentally validate the fibril formation rate expressions, we use data from two studies,^34,35^ where the formation and elongation of A *β* fibrils was probed by monitoring the increase in ThT fluorescence intensity as a function of time. Experimental evidence suggests that A*β* peptide fibrillation follows a template-dependent dock–lock mechanism.^5^ Typical ThT assays are bulk aggregation monitoring experiments, where the ThT fluorescence is proportional to the amount of formed amyloid fibrils, but the technique is not specific for individual fibrils. ThT fluorescence monitors fibrils of various sizes, oligomers, and other small aggregates. ThT fluorescence increases as the fibrillation reaction progresses because the *β*-sheet structures increase proportionately to the increasing length of fibrils. The observable fluorescence is related to protein concentration by Beer’s Law. For the case of fibril elongation, the observable fluorescence for formation of fibrils will follow the relationship:

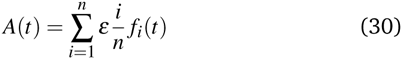

where *A*(*t*) is the cumulative absorbance at time *t*, *f*_*i*_(*t*) is the concentration of fibril of size *i* at time *t*, and *ε* is a scaling factor. Note that we do not include the intermediate “docked” monomer-template fibril complex in the above expression, because they are short-lived. To model this expression, We substitute the numerical solution of the ordinary differential equation system, eqns (4)–(8), into the fibril concentration term and calculate the weighted sum of the observable species.

The observable eqn (30) is validated by how accurately it can monitor an increase in fluorescence, which is a measure of the formation of fibrils as a function of time. Xiao et al.^35^ studies A*β*(1-40) elongation by monitoring docking and locking of monomers to A*β*(1-40) fibrils seeded with A*β*(1-40) template fibril. The ThT time course of A*β*(1-40) fibrillation shows an excellent fit to the observable eqn (30) using the MATLAB^®^ R2016a optimization routine lsqnonlin (see **Fig. 5A**). Dyes and extrinsic fluorescent probe bulk assays of protein elongation in the presence of seed fibrils do not exhibit the characteristic sigmoidal curve and generally lack an “observable lag time phase.”^6^ As expected, both Xiao et al. ^35^ experimental data and our model dynamics do not exhibit the characteristic sigmoidal curve, as the experimental assays is seeded with template fibrils. Narayan et al.^34^ also explore the kinetics of A*β*(1-40) aggregation by monitoring it through a ThT assay. A*β*(1-40) elongation follows a a hyperbolic curve without a detectable lag phase upon agitation. **Fig. 5B** compares the fitted observable eqn (30) with the Narayan et al.^34^ ThT assay of A *β*(1-40) elongation.

**Fig. 5.**
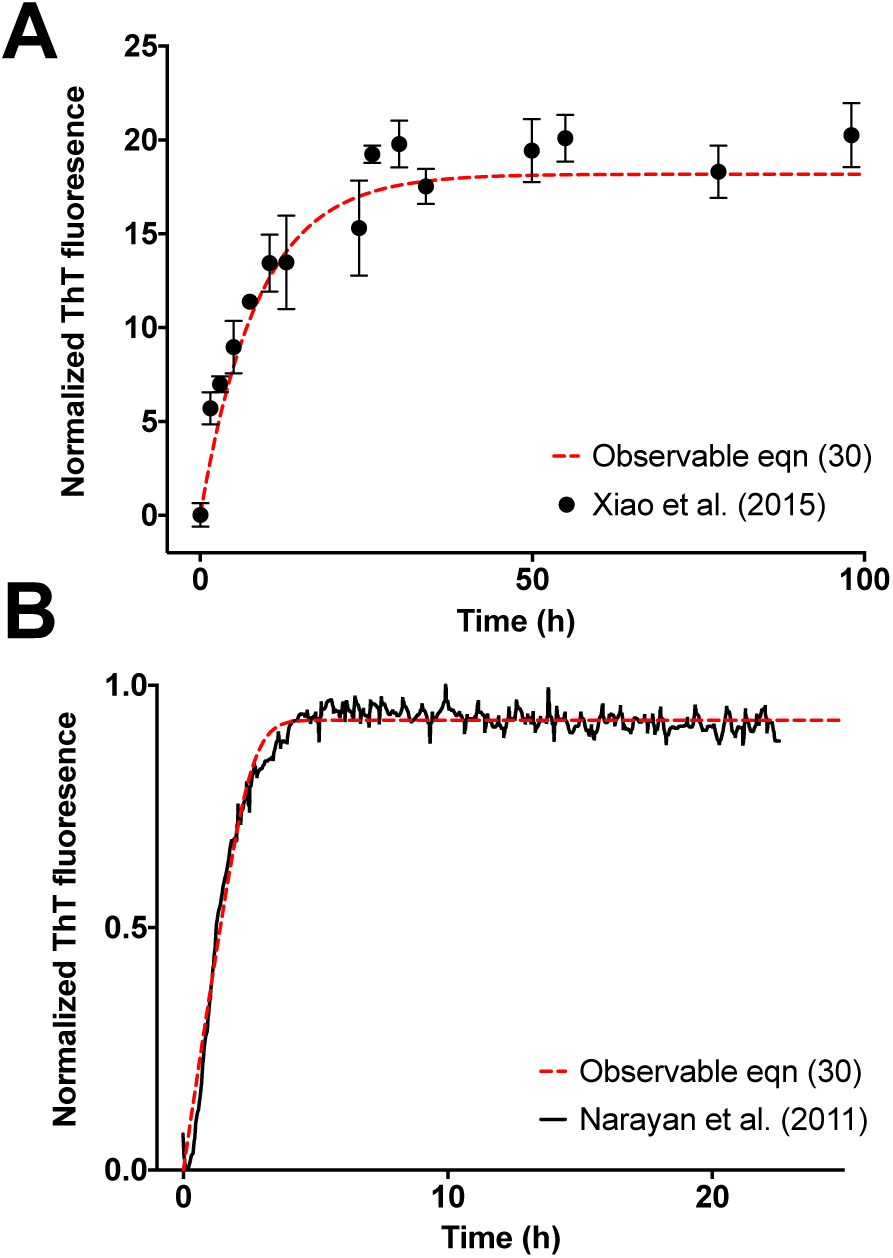
Validation of observable rate expression for fibril elongation using ThT assay experimental data for A *β* elongation. The observable eqn (30) governs well the increase in fluorescence intensity as function of time. Kinetic parameters were estimated with the nonlinear least squares optimization routine `lsqnonlin` from MATLAB^®^ R2016a. (A)Elongation of A *β*(1-40) incubated with A *β*(1-40) seed fibrils from **Fig. 4** of Xiao et al.^35^. The non-linear least-square estimation of kinetic parameter is: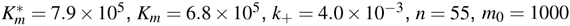, and 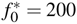. (B) Elongation of A*β*(1-40) incubated with seed fibrils from Fig. 1A of Narayan et al.^34^ The non-linear least-square estimation of kinetic parameter is: 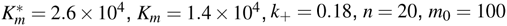, and 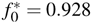 Experimental data was extracted with the digitizing software `im2graph` (http://im2graph.co.il).

## 7 Discussion

We have shown that it is possible to derive a lag time expression as a function of the kinetic parameters by writing the rate equations governing the reaction mechanism. In this work, we use the subsequent monomer addition dock-lock growth reaction mechanism (3) as a case study. For this reaction mechanism, we define the time lag for a final fibril, *F*_*n*_, as the sum of all of the timescales for the intermediate “locked" fibril species leading to the formation of the final fibril. This expression is a result of the causal connectivity in the subsequent monomer addition model, in which a monomer is docked and locked onto each intermediate, delaying the formation of the final fibril. The experimental implementation of our kinetic model for lag time requires a care-ful choice of initial conditions to satisfy the reactant stationary approximation and excess monomer condition, along with a careful consideration of the mechanism of interest.

In our subsequent monomer addition model, fibril elongation is nucleation-independent. Many current nucleation-dependent polymerization models rely on high order kinetics and unlikely collisions between multiple monomers to create a nucleus.^12,15,36^ It is widely believed that the time lag is a result of the nucleation process.^6,12^ Interestingly, we found that the characteristic ‘S’ shape sigmoidal curve typical of a lag time fibrillation process can be observed in a single-molecule study independent of primary and secondary nucleation processes. This finding is in agreement with earlier work by Rangarajan and de Levie^37^ and Baldassarre et al.^38^ Monitoring single-molecule species, we observe a lag phase in the absence of nucleation, which shows that the lag time is not necessarily a ‘wait time’ for the nucleus to form or the result of secondary nucleation. Though elongation is sufficient to produce a lag phase at the single-molecule level, it is not necessarily the sole contributor to the lag time. The contributions of nucleation and elongation to the lag time should be further explored by investigating other reaction mechanisms and implementing advanced single-molecule techniques experimentally.

The duration of the lag time exhibits a nonlinear relationship as a function of the kinetic parameters for the subsequent monomer dock-lock growth reaction mechanism. In the first association-dissociation reaction, the equilibrium constant depends nonlinearly on the concentrations of the monomer, fibril, and intermediate complex. Therefore, small changes to the concentrations will trigger larger changes in the equilibrium constant, and consequently, in the lag time. For example, the lag time increases exponentially with increases to the ratio between the dissociation constant of the monomer from the complex *K*_*m*_ and the apparent dissociation constant of the monomer from the complex 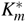, while the lag time decreases exponentially with increases to the on-rate constant, *k*_+_ (see **Fig. 4B**–C).

We found that the lag time does not change significantly with increasing concentration of the fibril template 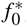. If a nucleation process is associated to our reaction mechanism, the nucleating species could be analogous to the fibril template species. Several studies suggest that increasing the concentration of seed available (via secondary nucleation, sonication-induced fragmentation, or increased initial concentration) decreases the lag time of fibrillogenesis.^39–41^ However, in our case, there is a slight increase in the lag time with increases in the ratio of 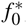 to *m*_0_ when *m*_0_ is in excess. The observed increase in lag time can likely be attributed to differences in the nature of dock-lock reaction mechanism (3). In our model, the fibril templates are depleted and not replenished to remain at a steady-state value. The addition of a nucleation process will change the derivation of the lag time.

Our analysis illustrates that there are important differences in the temporal dynamics of fibril elongation, which depend on the analytical technique used to monitor the total amount of aggregated proteins. Fibril elongation does not exhibit an observable sigmoidal-shape and lag phase when monitored with dyes and extrinsic fluorescent probes bulk assays, such as ThT fluorescence intensity experiments. However, this does not imply that there is no lag time associated with fibril elongation at the singlemolecule level. In our model, the concentration of the final fibril displays a lag phase. However, when considering ThT fluorescence intensity, which measures the amount of multiple aggregates and fibril species, the lag phase of the final fibril is hidden.

Our approach to derive a mathematical expression for the lag time could be used to investigate the factors affecting the length of the lag time for reaction mechanisms with off-path aggregation, inhibition kinetics, fragmentation, higher-order oligomerization, or other alternative pathways. The Knowles et al.^15^ model provides a more complete description of the protein aggregation process, including nucleation, elongation, and secondary nucleation or fragmentation. Powers and Powers^42,43^ also implements a nucleation-dependent polymerization mechanism with off-pathway aggregation, in which forming a nucleus is energetically unfavorable, but serves as a template for elongation. Furthermore, although the kinetics of amyloid formation are well-studied across many different peptides, alternative mechanisms are continually proposed and the process is far from being completely understood. Regardless of the mechanism of interest, applying our protocol for the calculation of timescales outlined to proposed mechanisms helps to elucidate the meaning of the lag time in single-molecule and bulk experimental studies. Since our unique expressions for the timescales of each reacting species are dependent on the kinetic parameters for a reaction mechanism, our protocol for estimating a lag time will serve as a basis for understanding the factors affecting complex protein aggregation and polymerization processes.

## Acknowledgements

This work is supported by the University of Michigan Protein Folding Diseases Initiative.

